# Comparative metagenomic analysis of diarrheal and non-diarrheal gut microbiome delineating the prospective development of prognostic markers and probiotics to protect from diarrhea

**DOI:** 10.1101/2023.07.17.549426

**Authors:** Rituparna De, Suman Kanungo, Asish K. Mukhopadhyay, Shanta Dutta

## Abstract

A cross-sectional gut microbiome analysis of 23 non-diarrheal and 5 diarrheal fecal samples was conducted by employing 16s rRNA amplicon sequencing and subsequent analysis for taxonomic profiling of OTUs and abundance interpretation of reads. Significant differences in the structural composition of the two groups were observed. In both Firmicutes was the most abundant phylum in majority of the samples. B/F ratio was consistently <1 in all diarrheal samples. Significant difference in mean B/F ratio of the two groups was found. Proteobacteria was significantly more abundant in the diarrheal group. *Prevotellaceae* was the most abundant family in non-diarrheal samples and was suppressed significantly in diarrheal samples. *Streptococcaceae* was the most abundant family in 60% diarrheal samples and where *Streptococcaceae* was suppressed, *Bacteroideaceae* and *Nocardeaceae* were the most abundant. In non-diarrheal samples where *Streptococcaceae* was almost completely suppressed *Bifidobacteriaceae* was the most abundant and suppressed other families significantly. A negative correlation was observed between *Prevotellaceae* and *Bacteroideaceae* in the non-diarrheal group. *Prevotella copri* was the most abundant species in 70% non-diarrheal samples and was significantly suppressed in diarrheal samples. *Proteus mirabilis* was identified in all the non-diarrheal samples while they were absent in diarrheal samples. The OTUs associated with diarrheal dysbiosis can serve as prognostic markers. This is the first report on the comparative analysis of diarrheal and non-diarrheal microbiome, to our knowledge, and distinctly addressing the gut microbiome dysbiosis from the context that can lead to the development of prognostic markers and probiotics for protecting the endemic population from diarrhea.

## Introduction

Diarrhea continues to be a major contributor of mortality, morbidity and a parameter of socio-economic loss globally (1). It occurs due to the lack of sanitation and is a recurring threat in the endemic regions (2). Advanced prognostic measures can help to avert diarrhea and can help to assuage disease burden.

Next-generation sequencing is being increasingly used as a supplementary tool to augment precise and rapid diagnosis alongside traditional phenotypic and genotypic methods in molecular epidemiology laboratories worldwide (3). With their immense potential to identify diverse pathogens simultaneously and also predict their abundance in clinical samples, they hold immense prospects of serving as novel prognostic methods based on the identification of distinct microbial taxa (4).

In order to construe the structural diversity and abundance of the various taxa comprising the diarrheal and non-diarrheal gut microbiome a comparative metagenomic analysis was undertaken on the Illumina MiSeq platform and OTU reads generated by 16S rRNA amplicon sequencing were matched to the gene database using the QIIME pipeline and identified. Population-based inferential statistical analysis of OTU abundance confirmed significant differences in composition of the gut microbiome in the two groups. This helped us to identify dysbiosis-associated organisms which are either enriched or suppressed in the diarrheal gut microbiome. These specific microbial signatures can be developed as prognostic markers to reduce diarrheal disease burden.

## Materials and Methods

23 non-diarrheal fecal samples (CS1-CS22 and CS25) from Kolkata and 1 sample each from 5 hospitalized patients of acute diarrhea (21-25) at the Infectious Diseases Hospital, Kolkata, were collected. DNA was extracted following the GITC method (4). The 16S rRNA gene V3-V4 regions were amplified (4) followed by addition of Illumina sequencing barcoded adaptors (Illumina, CA, USA.). The libraries were normalized and pooled for multiplex sequencing using the Illumina MiSeq v3 600 cycles cartridge (Illumina, CA, USA). The run was performed in paired end mode with 275 bp read length for each of forward and reverse reads.

The paired end reads were demultiplexed using bcl2fastq tool, quality checked using FastQC, filtered for high quality (Q30) reads using cutadapt program (5) and joined using Fastq-join (6). These were considered for microbiome search using the QIIME pipeline (7). The query sequences were clustered using the UCLUST (8) method against the Greengene database (9) and taxonomies were assigned at >=97% sequence similarity. Downstream analysis and visualization was performed using R-package (10). Relative abundance from phylum to species was calculated from read counts assigned to OTU divided by total utilized reads for microbiome search and was presented as stacked column plot. B/F ratio was calculated as the ratio of Bacteroidetes to Firmicutes in each sample. The difference in mean relative abundance was calculated for the major phyla and families for the two groups and the results were presented using stacked bar-diagram. The significance of this difference was calculated with students’ two-tailed T-test and correlation with the help of Pearson correlation coefficient. Z score was used to analyze difference of proportion of the two groups with respect to a taxon.

## Results

22 different phyla were identified. Actinobacteria, Proteobacteria, Firmicutes, Bacteroidetes were present in all the samples. The comparative relative abundance of the major phyla and their frequency of occurrence in the two groups have been presented in Figure 1. Firmicutes was the most dominant phylum in 80% diarrheal and 52.2% non-diarrheal samples with a mean abundance of 52% in diarrheal and 42% in non-diarrheal. Bacteroidetes were dominant in 43.5% non-diarrheal samples but in none of the diarrheal samples. The mean abundance of Bacteroidetes was 7.72% in diarrheal and 41.71% in non-diarrheal (control) samples. The difference was significant with *p*-value 0.000114. The average B/F ratio in diarrheal samples was 0.23 and was consistently <1 in all the diarrheal samples while in non-diarrheal samples the average was 1.23 with 11 out of 23 samples having a B/F ratio >1. B/F ratios is significantly different with a *p*-value of 0.008228. The mean abundance of Proteobacteria was 21.76% in diarrheal and only 3% in non-diarrheal samples and the difference was significant with *p*-value was < .00001. Analysis of correlation of abundance of Proteobacteria and Firmicutes showed that in the diarrheal samples r(3)= -0.6558 indicating moderate negative correlation while in the non-diarrheal samples r(21) = -0.4015 indicating weak negative correlation and the result was significant with p= .057914 at p < .10. Analysis of correlation of abundance of Proteobacteria and Bacteroidetes showed a weak negative correlation r(3)= -0.3082 in diarrheal samples and a weak positive correlation r(21)= 0.3656, was observed in the control. Phyla TM7 and Fusobacteria were significantly higher in diarrheal samples compared to non-diarrhealsamples (Table 1).

**Table 1.** showing the difference in abundance of major phyla and families in diarrheal and non-diarrheal groups.

| Phyla | Actinobacteria | Proteobacteria | Firmicutes | Bacteroidetes | Verrucomicrobia | Fusobacteria | TM7 | Tenericutes | Synergistetes |
| --- | --- | --- | --- | --- | --- | --- | --- | --- | --- |
| Diarrheal | 17.122 | 21.7584 | 51.97764 | 7.72244 | 0.65012 | 0.20918 | 0.20682 | 0.00868 | **0.00126** |
| Non-diarrheal | 12.124 | 3.01725652 | 42.21951 | 41.7104 | 0.6184304 | 0.01335652<br>2 | 0.02928 | 0.22083 | **0.0110043** |
| t-test | 0.5894 | 6.74676. | 1.17297 | -4.53844 | 0.02848 | 3.741 | 2.11332 | -0.93385 | **-0.69884** |
| p-value | 0.5606 | < .00001 | 0.251443 | 0.000114 | 0.977495 | 0.000915 | 0.04433 | 0.358973 | **0.490857** |
| Families | Lachnospiraceae | Lactobacillaceae | Erysipelotrichaceae | Prevotellaceae | Streptococcaceae | Enterobacteriaceae | Coriobacteriaceae | Veillonellaceae | Ruminococcaceae |
| Diarrheal | 1.682 | 0.28276 | 1.370 | 0.361 | 30.1609 | 9.6501 | 1.409 | 1.22158 | 8.38546 |
| Non-diarrheal | 5.724<br>7478<br>26 | 0.335078<br>261 | 6.567<br>1391<br>3 | 27.17<br>99 | 1.72402<br>6087 | 2.0259 | 4.643<br>8347<br>83 | 7.26865<br>217<br>4 | 17.36140<br>87 |
| t-test | -2.473 | -0.096 | -1.334 | -3.065 | 4.710 | 4.248 | -1.509 | -2.1869 | -1.72 |
| p- value | .010<br>signifi-<br>cant | .462303 | .0968<br>signifi-<br>cant | .00502<br>signifi-<br>cant | .000036<br>significa-<br>nt | .000122<br>significnt | .0716 | .01896<br>significa-<br>nt | .048693<br>significant |

**Figure 1.**
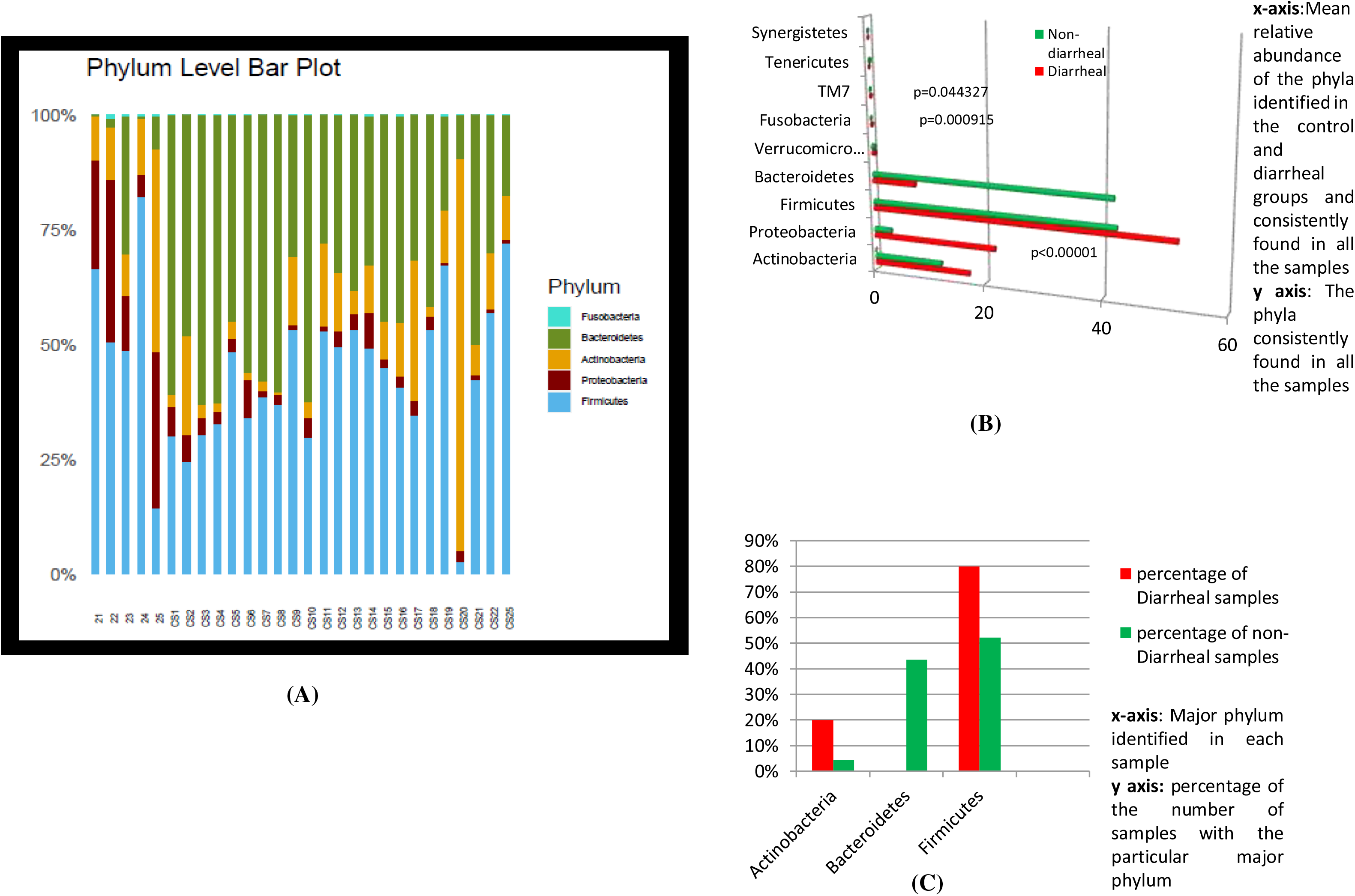
showing (A) the relative abundance of and (B) difference in relative abundance of the major phyla in 5 diarrheal samples 21-25 and non-diarrheal samples CS1-CS22 and CS25(C) shows the frequency of finding Actinobacteria, Firmicutes, Bacteroidetes as the most dominant phylum in the population expressed as percentage of diarrheal and non-diarrheal samples carrying the particular phylum

**Figure 2.**
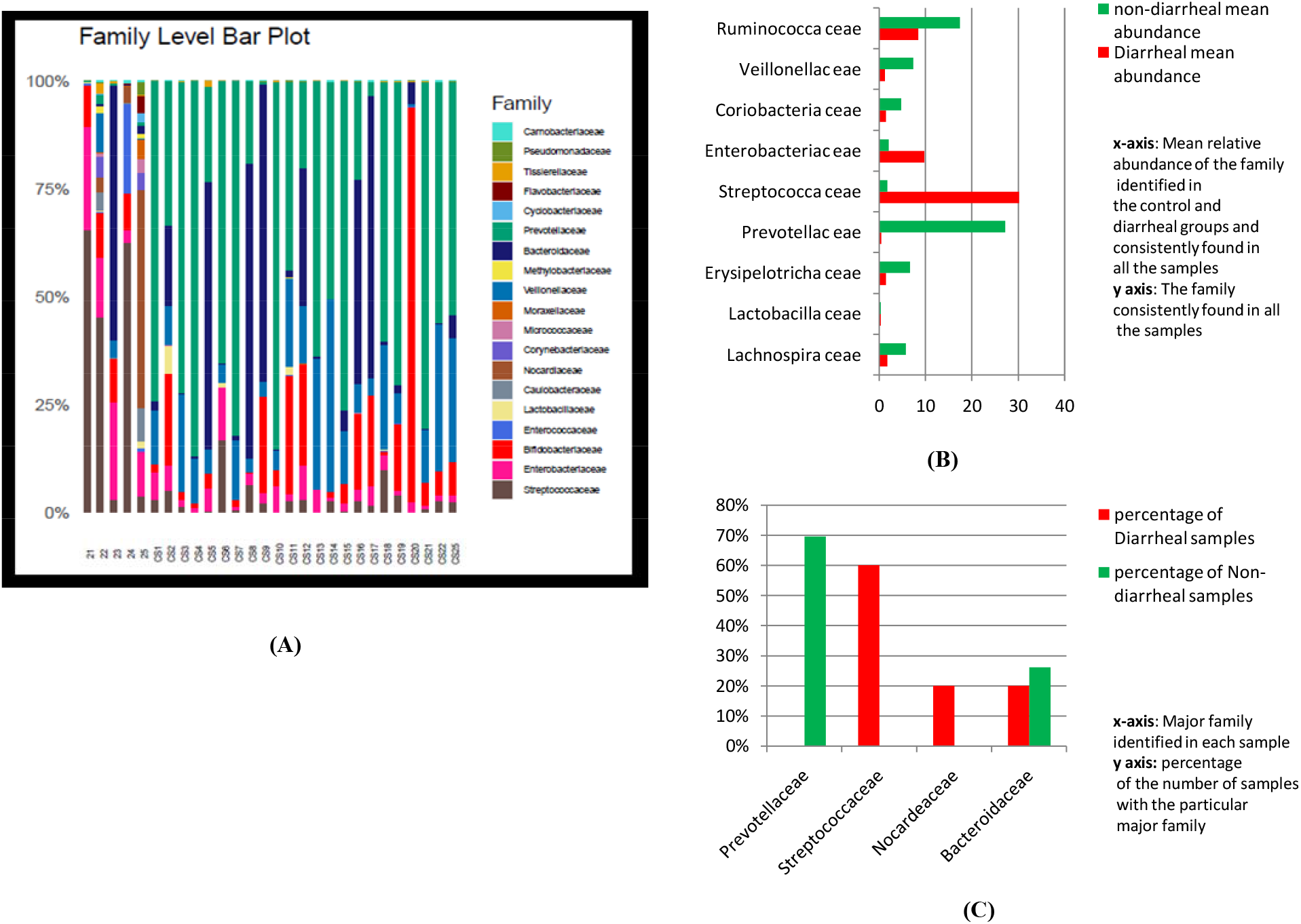
showing (A) the of the relative abundance of and (B) difference in relative abundance of the major families in 5 diarrheal samples 21-25 and non-diarrheal samples CS1-CS22 and CS25 (C) shows the frequency of finding *Prevotellaceae, Streptococcaceae, Nocardeaceae, Bacteroidace ae* as the most dominant phylum in the population expressed as percentage of diarrheal and non-diarrheal samples carrying the particular family

Significant differences in abundance was observed for *Lachnospiraceae, Erysipelotrichaceae, Prevotellaceae, Streptococcaceae, Enterobacteriaceae, Vellionellaceae* and *Ruminococcaceae. Prevotellaceae* was the most abundant family in 69.6% non-diarrheal samples and was suppressed significantly in diarrheal samples while *Streptococcaceae* was the most abundant family in 60% diarrheal samples. Both the differences in frequency of occurrence in the two groups expressed by Z-score was significant with p-values 0.004 and 0.00008 respectively. In diarrheal samples in which *Streptococcaceae* was suppressed, *Bacteroideaceae* and *Nocardeaceae* were the most abundant. In non-diarrheal samples where *Streptococcaceae* was almost completely suppressed *Bifidobacteriaceae* was the most abundant and suppressed other families significantly. A negative correlation was observed between *Prevotellaceae* and *Bacteroideaceae* in the control.

In diarrheal samples negative correlation of *Prevotellaceae* and *Lachnospiraceae* was found with *Streptococcaceae* and *Enterobacteriaceae* while *Ruminococcaceae* was found to have negative correlation with *Enterobacteriaceae*. In non-diarrheal samples significant negative correlation was found between *Lachnospiraceae* and *Enterobacteriaceae*. The significance of differences in relative abundance of the major phyla and families and their correlation have been presented in Tables 1 and 2.

**Table 2.**
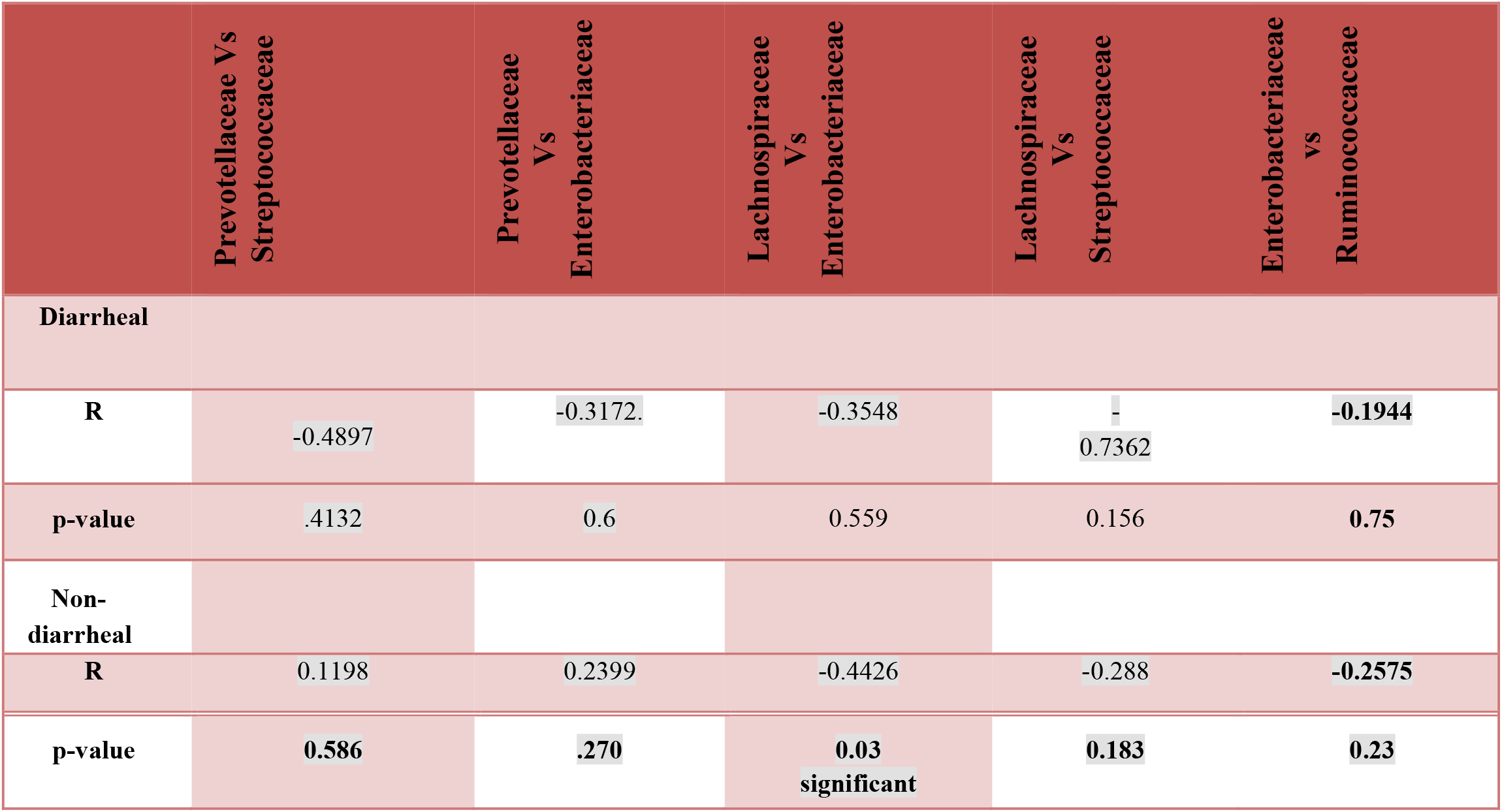
showing the correlation between different families in diarrheal and non-diarrheal groups.

*Proteus mirabilis* was found associated with all non-diarrheal samples while it was absent in diarrheal samples. *P. copri* was found in all the samples of both the groups though the mean abundance in diarrheal samples was 14.86% while in non-diarrheal samples was 22.21%. The difference was found to be significant with *p*-value .012063.

## Discussion

Previously we have catalogued and analyzed the significance of structural diversity in the diarrheal gut microbiome (5). It prompted our interest to scrutinize the community gut microbiome. Our analysis culminated in a hypothesis that the OTUs suppressed in the diarrheal microbiota and those enriched in the control may help in prognosis and drive the development of probiotics. The present study found a trend of negative correlation of commensals and pathobionts among the 5 diarrheal samples we tested and is in complete accordance with our previous study (4). In the non-diarrheal microbiota we found encouraging contrasting trends of significant negative correlation between commensal *Lachnospiraceae* and pathobiont *Enterobacteriaceae*. This could serve as a prognostic marker for screening vulnerability to diarrhea. *Bifidobacteriaceae* was found to be dominant over pathobionts. Also, family *Prevotellaceae* that was significantly enriched in controls and the presence of significantly higher abundance of *P. copri* compared to cases can also serve as a marker of diarrheal dysbiosis. The findings were reinforced by epidemiological data that the frequency of occurrence of these markers were significantly different in the two groups. These may be developed as probiotics for the endemic population to promote normobiosis and prevent diarrhea. *P. copri* has been already found to be a potential next-generation probiotic (11).

Another interesting feature revealed by our study was the total absence of *P. mirabilis* in diarrheal patients. In our previous study it was absent in all but one in which its relative abundance was 0.00025% (4).

The present study is the first report on the comparative analysis of diarrheal and non-diarrheal gut microbiome. It identified prospective OTUs which can serve as potential prognostic markers and can be developed to provide protection among endemic population, particularly, in the economically backward areas of the world where gut microbiome dysbiosis due to different parameters like malnourishment can make them vulnerable to diarrheal pathogens (12).

## Importance

This is the first report of a comparative analysis of gut microbiome of diarrheal and non-diarrheal fecal samples. It presents the structural dysbiosis of the diarrheal gut microbiome compared to the non-diarrheal community counterpart. It helped in the identification of prospective commensal candidates that may be useful for counteracting the pathobionts for normobiosis. Commensals were significantly enriched in the non-diarrheal subjects while they were significantly suppressed in diarrheal patients. A clear association of their beneficial effect was visible upon overriding the pathobiont communities and consequently shaping the gut microbiota. These are potential markers of prognosis and can be developed into probiotics for providing protection to population vulnerable to diarrheal pathogens, specially, in the endemic regions of the world. This will help to implement these as routine formulations to revert the gut microbiome to a healthy condition to minimize dysbiosis and help reduce diarrheal disease burden.

## Availability of data

The study has been registered in the European Nucleotide Archive (ENA) at EMBL-EBI under accession number ERC000029 and sequences will be deposited.

## Data reporting

The study is being reported following STORMS (supplementary file).

## Acknowledgement

RD is grateful to Dr. G. B. Nair for his help in this project and inspiration for microbiome research.

